# Unravelling the molecular mechanisms of fish salinity adaptation in the face of multiple stressors: A comparative multi-tissue transcriptomic study in the Llobregat River, Barcelona, Spain

**DOI:** 10.1101/2023.11.20.567554

**Authors:** Camilo Escobar-Sierra, Miguel Cañedo-Argüelles, Dolors Vinyoles, Kathrin P. Lampert

## Abstract

Freshwater salinization poses a growing global environmental concern, introducing complex chemical cocktails and jeopardizing freshwater biodiversity, particularly fish populations. This research aimed to elucidate the molecular foundations of salinity adaptation in a non-native minnow species (*Phoxinus septimaniae* x *P. dragarum*) exposed to saline effluents from potash mines in the Llobregat River, Barcelona, Spain. Employing high-throughput mRNA sequencing and differential gene expression analyses, we examined brain, gills, and liver tissues collected from fish at two stations (upstream and downstream of saline effluent discharge). Salinization markedly influenced global gene expression profiles, with the brain exhibiting the most differentially expressed genes, emphasizing its unique sensitivity to salinity fluctuations. Pathway analyses revealed the expected enrichment of ion transport and osmoregulation pathways across all tissues. Furthermore, tissue-specific pathways associated with stress, reproduction, growth, immune responses, methylation, and neurological development were identified in the context of salinization. Rigorous validation of RNA-seq data through quantitative PCR (qPCR) underscored the robustness and consistency of our findings across platforms. This investigation unveils intricate molecular mechanisms steering salinity adaptation in non-native minnows confronting diverse environmental stressors. Advancing our comprehension of genomic responses to salinity changes, our study provides crucial insights into the adaptive strategies of aquatic organisms grappling with freshwater salinization. This comprehensive analysis sheds light on the underlying genetic and physiological mechanisms governing fish adaptation in salinity-stressed environments, offering essential knowledge for the conservation and management of freshwater ecosystems facing escalating salinization pressures.

## 1. Introduction

Human activities such as transportation (Hintz & Relyea, 2019), resource extraction (Cañedo-Argüelles et al., 2013) and agriculture (Thorslund et al., 2021) are leading to an increase in the salinity of freshwater ecosystems around the world (i.e. freshwater salinization). This is accompanied by other pollution sources (e.g. agricultural activities not only lead to salinization but also to fertilizer and pesticide pollution) that lead to novel chemical cocktails (Kaushal et al., 2019). Since freshwater organisms can only tolerate certain salinity ranges, freshwater salinization is leading to drastic reductions in aquatic biodiversity (Cañedo-Argüelles et al., 2019). However, our understanding of the ecological impacts of freshwater salinization is still limited. While fish have been identified as potentially affected by salinization, they have received considerably less attention than other freshwater organisms (Cunillera-Montcusí et al., 2022).

Potash mining operations often store their wastes in open areas near the mines, creating artificial salt mountains, primarily composed of NaCl, such as the case of our study site the "Potasses del Llobregat" potash mine situated near the Llobregat river in Barcelona, Spain. These mountains of salt-rich waste have the potential to generate highly saline effluents that are discharged into surrounding rivers and streams, becoming the main driver of freshwater salinization in certain areas (Otero & Soler, 2002). Although brines are usually collected and treated and/or transported to the sea, many technical problems persist (e.g. leaks in the collection systems, saline water infiltration) that can lead to severe salinization (Gorostiza et al., 2022; Gorostiza & Saurí, 2019). For example, in the Soldevila stream (NE Spain, very close to our study site) the electrical conductivity was 132.4 mS/cm (Ladrera et al., 2016), much higher than that of the sea. These point source saline pollution effluents associated with potash mining activities are known to severely reduce the diversity of native freshwater macroinvertebrate species (Bäthe & Coring, 2011; Cañedo-Argüelles et al., 2017) and promote invasions (Arle & Wagner, 2013; Braukmann & Böhme, 2011; Lewin et al., 2018), but the effect on fish communities remains largely unknown.

Urban-influenced rivers are also affected by nonpoint source pollution and climate variability, which give rise to novel elemental combinations, referred to as ’chemical cocktails’ in watersheds, acting as multiple stressors on stream fauna (Kaushal et al., 2019; Schäfer et al., 2023). Consequently, relying solely on traditional methods to evaluate the effects of multiple stressors on factors such as endpoint mortality, or population and community, might be limiting. Novel high-throughput methods such as transcriptomics have the potential to assess the entire gene expression of organisms exposed to multiple stressors in the wild (Lowe et al., 2017). Furthermore, transcriptomics can provide insight in the physiological status of species by measuring responses to multiple stressors that may or may not be known in advance of sampling (Jeffries et al., 2021). Consequently, utilizing transcriptomic methods can aid in evaluating the physiological responses of wild organisms, potentially informing management decisions (Connon et al., 2018). This method has demonstrated success in prior research involving wild fish populations, effectively identifying the stressors influencing fish physiological responses amidst varying environmental conditions, with salinity acting as one of the main stressors (Escobar-Sierra & Lampert, 2023; Jeffrey et al., 2023; Komoroske et al., 2016). This gene expression data obtained under natural conditions, can help elucidate the affected molecular pathways and identify the stressors within these complex mixtures that impact organisms.

Although our study site, the Llobregat River, experiences multiple stressors, freshwater salinization from potash mining stands out as the dominant stressor affecting aquatic fauna. The hyperosmotic stress poses significant physiological challenges to many freshwater animals, especially fish, impacting their osmoregulation processes (Guh et al., 2015). The gill, as a primary organ for osmotic homeostasis, has been the focus of osmotic stress transcriptomic research, revealing differential expression of genes associated with immune system, membrane transport, and regulation pathways in diverse fish species exposed to varying salinities (X. Chen et al., 2021; Cui et al., 2019; Escobar-Sierra & Lampert, 2023; Guo et al., 2018a; Lee et al., 2020; Su et al., 2020a). Recent studies have also identified differential gene expression in the brain and liver of certain fish species under osmotic stress, emphasizing pathways related to immunity, reproduction, growth, osmoregulation, energy metabolism, oxidative stress, and ion transporters (Lin et al., 2020; Liu et al., 2018; K. Zhou et al., 2020). Even though for specific tissues transcriptomic resources in fish exist, comprehensive studies assessing multiple tissue-specific salinity responses in fish remain limited. This study aims to address this gap by conducting a comparative transcriptomic analysis focusing on gill, brain, and liver tissues under salinity stress.

The genus *Phoxinus*, presents a diverse array of species distributed across the northern hemisphere, particularly in Europe and parts of northern Asia (Palandačić et al., 2020). Known for their adaptability, these small freshwater fish populate various habitats from mountain streams to lowland rivers and lakes (Frost, 1943). Several *Phoxinus* spp. species have been introduced to new catchments, notably in the Llobregat basin an introduction event has created a hybrid species resulting from the translocation of *P. septimaniae* from southern France and *P. dragarum* from the Garonne River (Corral[Lou et al., 2019). The *Phoxinus spp.* invasions into several new catchments have led to ecological consequences, such as competition for food and predation on eggs, adversely affecting native species (Borgstrøm et al., 2010; Garcia-Raventós et al., 2020; Tiberti et al., 2019, 2022). Notably, their adaptability to high salinity in brackish waters, observed in certain coastal regions, may bolster their potential to invade upstream and colonize systems facing altered ecological conditions due to freshwater salinization. Similar adaptive advantages in increasing salinity scenarios have been described in other freshwater species, leading to expanded populations and distributions (Dobrzycka-Krahel & Fidalgo, 2023; Hudson et al., 2021; Olin et al., 2022; Piscart et al., 2011). Although *Phoxinus* spp. euryhalinity suggests they have the advantage to tolerate the salinity changes, it has been estimated that the activation of the hyperosmotic osmoregulatory mechanisms for euryhaline fishes can account for 20-68% of their total energy expenditure (Bœuf & Payan, 2001; Komoroske et al., 2016). These costs may drive the need for physiological and behavioural optimization to minimize energy expenditure, resulting in narrow physiological tolerance windows and behavioural avoidance of salinities outside the optimal range (Dowd et al., 2010; Komoroske et al., 2016). However, there is still a lack of knowledge on the molecular mechanisms underlying this species response to salinity stress. Their invasive potential, salinization adaptability and wide distribution make the minnow a valuable model to study the osmoregulation mechanisms of euryhaline species under changing salinity concentrations.

The primary objectives of this study encompass understanding the molecular underpinnings of salinity adaptation in *Phoxinus septimaniae* x *P. dragarum* (from now on minnow) populations inhabiting contrasting salinity conditions and multiple stressors related to the potash mining activity in the Llobregat River in Barcelona, Spain. Our research seeks to unravel the specific effects of freshwater salinization on the gene expression profiles of brain, gills, and liver tissues, targeting the identification of tissue-specific physiological pathways amidst heightened salinity levels. We hypothesized that fish inhabiting high-salinity environments should exhibit distinct and tissue-specific gene expression patterns related to adaptive mechanisms in the face of increased salinity. To address this hypothesis, high-throughput mRNA sequencing and differential gene expression analyses were employed to explore the gene expression profiles of minnow populations at different salinity stations. Additionally, the proposed methods include conducting detailed transcriptomic analyses and differential gene expression studies to decode the tissue-specific responses and shared biological processes vital for the adaptation of aquatic organisms facing rising salinization pressures. These analytical techniques allow for the identification of key genetic and physiological mechanisms essential for adaptation in salinity-stressed environments.

## 1. Material and methods

### 1.1. Sampling design

We selected two sampling stations with distinct salinity levels along the main channel of the Llobregat River in Barcelona, Spain for fish sampling on June 9, 2022. The salt-polluted station (from now on “Salt” in the figures) is located near the outflow of the "Potasses del Llobregat" potash mine, which continuously releases high salinity waste. In contrast, the "reference station" is approximately five kilometers upstream from the Salt-polluted station, situated near the Balsareny locality **(Figure 1).** At the time of sampling, the Control station had a water conductivity of 834μS, a salinity of 0.4ppt, and a temperature of 17.5°C. These are normal values for natural Mediterranean rivers and streams (Sánchez-Montoya et al., 2012), which have relatively high salt concentrations due to rock dissolution. In contrast, the Salt station exhibited a conductivity of 2641000μS, a salinity of 18.1ppt, and a temperature of 17.2°C. The difference in salinity between the two stations is staggering, with the Salt-polluted station being 45 times saltier than the reference station.

**Figure 1.**
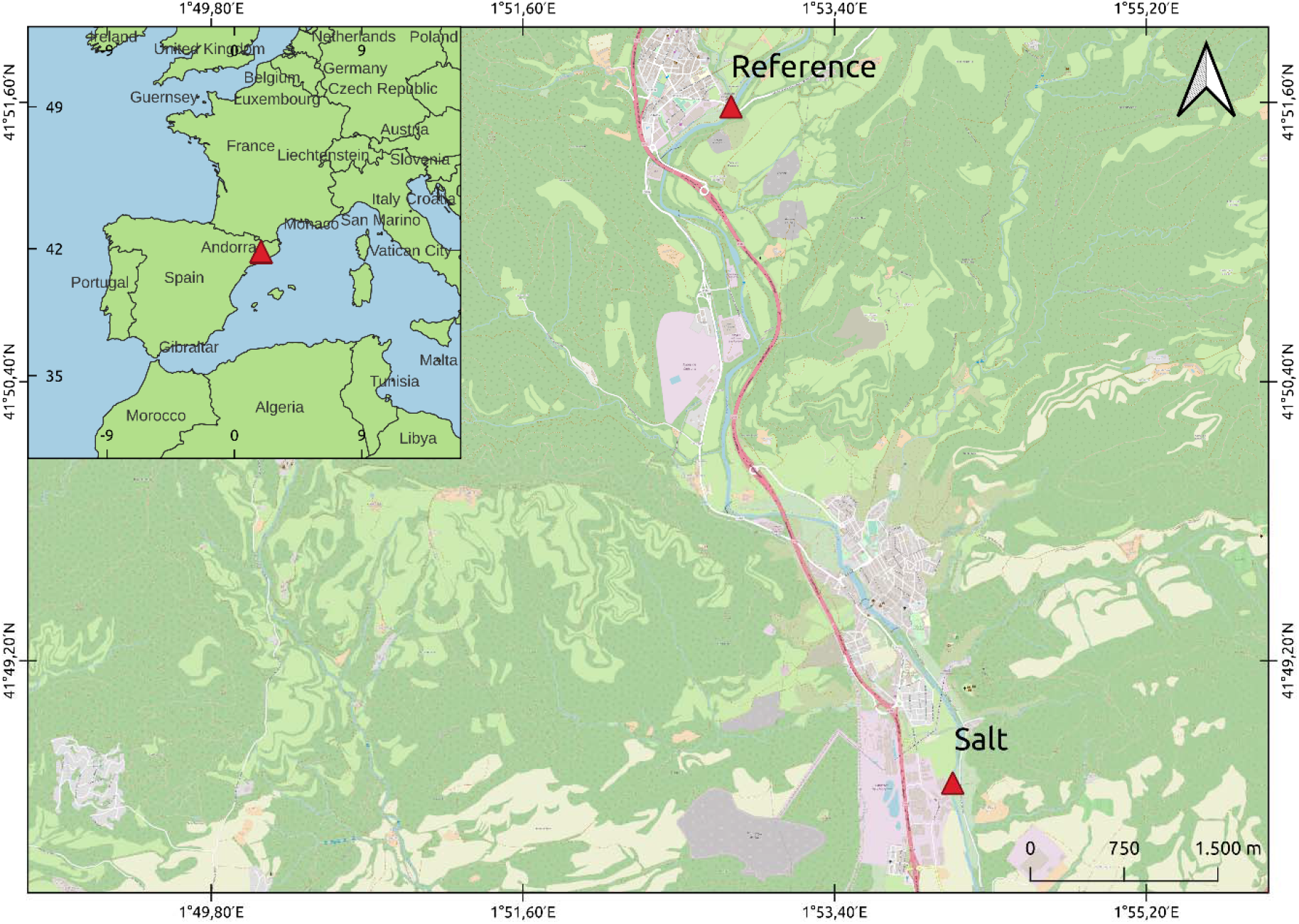
Locations of fish sampling stations in the Llobregat River catchment. Three fish were collected from each station, and brain, gill, and liver tissues were collected for RNAseq. Salinity at the salt-polluted station “Salt”: 18.1 ppt, and at the “Reference” station: 0.4 ppt.

Minnows were sampled by electrofishing using a portable unit which generated up to 200 V and 3 A pulsed direct current (DC) in an upstream direction. The fish collection was approved by the Regional Government of Catalonia (Ref. AP/004). All procedures were conducted in accordance with the European Directive for animal experimentation (2010/63/EU). After capture, fish were promptly euthanized with an overdose of tricaine methanesulfonate (MS-222 at 1g./L) followed by brain, liver, and gill tissue collection. The collected tissues were immediately fixed in RNAprotect (QIAGEN) and stored at -20°C. In total, we obtained six fish samples, with three from the Salt station and three from the reference station.

### 1.2. RNA isolation and Illumina sequencing

To isolate RNA, we homogenized the fixed tissue of each sample in a solution of 700μl RLT buffer, with a 10μl:1ml concentration of β-mercaptoethanol, using a FastPrep-24™ (MP Biomedicals) bead beater for 30 seconds at 5 m/s. Subsequently, we extracted the RNA using an RNeasy Mini Kit (QIAGEN), following the manufacturer’s instructions. We confirmed the quality of the RNA extractions using a Nanodrop 1000 Spectrophotometer (Peqlab Biotechnologie, Germany), ensuring a concentration >50ng/μl, an OD 260/280 ratio of 1.8-2.1, and an OD 260/280 ratio >1.5. The RNA Integrity Number (RINe) value was assessed using the RNA ScreenTape system in an Agilent 2200 TapeStation, confirming that the samples had a RINe score greater than 7.0. Subsequently, we used an Illumina Tru-Seq™ RNA Sample Preparation Kit (Illumina, San Diego, CA, USA) to generate sequencing libraries, following the manufacturer’s protocol. After purification, Illumina sequencing was performed on an Illumina/Hiseq-2500 platform (Illumina, San Diego, CA, USA) to generate 100 bp paired-end reads with a read depth of 50 million reads, as recommended by the Cologne Center for Genomics (CCG), Germany. The choice of read depth followed CCG technical advice, which estimated that this read depth would be sufficient for constructing a de novo transcriptome with eighteen biological replicates.

### 1.3. Sequence quality control, de novo assembly, and annotation

We conducted quality control checks on the raw sequences of all 18 samples using FASTQC (Andrews, 2010). Subsequently, we improved the quality by filtering out adaptor sequences and low-quality reads using TRIMMOMATIC, with a quality threshold of phred + 33, ensuring a minimum quality score of 25 for all bases, and retaining reads with a minimum length of 50 bp for downstream analyses (Bolger et al., 2014). We used TRINITY v2.9.1 (Grabherr et al., 2011) to create a de novo assembly with the nine pairs of clean sequences. The reads from the raw nine paired sequences were mapped to the de novo transcriptome using SALMON (Patro et al., 2017) as the abundance estimation method. To assess the completeness of the transcriptome assembly, we used BUSCO v5.2.2 (Simão et al., 2015) by searching against the vertebrata_odb10 dataset (Creation date: 2021-02-19). The expression matrix was generated, and low-expression transcripts (minimum expression of 1.0) were filtered using TRINITY v2.9.1, followed by the normalization of expression levels as transcripts per million transcripts (TPM). The identification of likely protein-coding regions in transcripts was performed using TRANSDECODER v5.5.0 (Haas et al., 2013). The filtered transcriptome sequencing reads were aligned to protein, signal peptides, and transmembrane domains using DIAMOND v2.0.8 (Buchfink et al., 2015), SIGNALP 6.0 (Teufel et al., 2022), TMHMM v2.0 (Krogh et al., 2001), and HMMER v3.3.2 (Finn et al., 2011). We functionally annotated the de novo transcriptome using TRINOTATE v3.2.2 (Grabherr et al., 2011).

### 1.4. Differential gene expression analysis

Prior to conducting differential gene expression analysis, we used the SALMON tool to align the clean reads of the three sites to the filtered transcriptome and calculated the mapping rate. The mapping tables were merged based on the stations and normalized using TMM (Robinson & Oshlack, 2010). The differential expression analysis, with three biological replicates for the three types of tissues, was performed to compare the Salt and Control stations using the DESeq2 package (Love et al., 2014). We set a fold change cutoff at >2 and an FDR≤0.05. This rigorous analysis allowed us to identify thousands of gene expression differences between the tissues at the control and salt stations. The incorporation of the false discovery rate (FDR) within DESeq2 is essential to control type I errors, making it a widely accepted standard in genomics for correcting false positive results in differential expression analysis (Benjamini & Hochberg, 1995; J. J. Chen et al., 2010). We visualized the differentially expressed genes (DEGs) using a heatmap plot created with GGPLOT2 (Wickham, 2016), highlighting the changes in expression levels for each tissue type at both stations. All the significant DEGs were classified according to their higher-level Gene Ontology (GO) terms. Subsequently, we conducted a GO pathway enrichment analysis using ShinyGo (Ge et al., 2020) (version 0.8) to identify GO terms with unbalanced distributions between the two groups of genes or probe sets. To account for multiple testing issues, we applied an FDR correction to the P-values in this comparison to control falsely rejected hypotheses (Khatri et al., 2012). We graphically represented the top pathways identified in both the higher GO terms DEGs classification and the enrichment analysis, focusing on their relevance to osmoregulation challenges based on previous studies. The assembly was performed in the High-performance computing system of the University of Cologne (CHEOPS), and the rest of the analysis was carried out using the Galaxy project platform (The Galaxy Community et al., 2022).

### 1.5. RNA-seq qPCR validation

To validate the RNA-seq results, we selected a subset of twenty-four genes from the DEGs dataset (Table S3) and assessed their expression using real-time quantitative PCR analysis (qPCR). The sequences were retrieved from the assembled transcriptome and used to design specific oligonucleotides with the Primer-Blast software (Ye et al., 2012). First, reverse transcription of the total RNA samples was performed using the RevertAid first-strand cDNA synthesis kit (Thermo Scientific), and the resulting cDNA was diluted tenfold for qPCR. The qPCR was performed using PowerUp™ SYBR™ Green Master Mix in a StepOne Real-Time PCR system following the manufacturer’s indications. Each gene was assessed using the 2^-△△Ct^ method, with ACTB as the housekeeping gene for normalization. We expressed the qPCR results as log_2_ fold changes and conducted a Spearman’s correlation analysis across all the samples to compare the expression values obtained from the RNAseq analysis.

## 2. Results

### 2.1. Sequence quality control, de novo assembly, and annotation

The RNA sequencing conducted on brain tissues yielded an average of 26.22 Mbp (± 2.14 Mbp SD) total transcripts after quality control. For gill tissues, the total transcripts averaged 19.77 Mbp (± 4.29 Mbp SD). Liver tissues had an average of 31.22 Mbp (± 6.09 Mbp SD) total transcripts. Notably, all retained reads after quality control had a quality score %Q30 of 100%. The assembly with all sequences produced a raw transcriptome of 708.3 mb with an N50 of 1579 bp. The number of putative genes for the Trinity assembly was 434823, with a total transcript number of 760990. We identified 3354 complete and fragmented BUSCOs with an annotation completeness of 97.3%. Following the filtration of low-expression transcripts, we retained 35.62% of the initial raw transcriptome, resulting in a filtered transcriptome of 280.1 mb, encompassing 155426 putative genes with an N50 of 1930 bp. Notably, 3354 complete BUSCOs were retained, with an assembly completeness of 66.00%. Out of the de novo transcriptome, 90104 transcripts were annotated using TRINOTATE, representing 42.03% of the total transcriptome.

### 2.2. Differential gene expression analysis

The analysis revealed considerable alterations in global gene transcription profiles induced by changes in salinity across the brain, gills, and liver tissues of the minnow. The average percentage of reads mapping as proper pairs was 48.82% (± 1.43% SD) for the brain, 48.95% (± 1.16% SD) for gills, and 64.35% (± 4.67% SD) for liver tissues. The principal component analysis (PCA) indicated tissue-specific expression patterns, showing that samples across both stations clustered together by tissue type **(Figure 2A).** Along the PC1 (70.78% variance explained), the sequences were segregated based on tissue type, clearly delineating between brain, gills, and liver tissues. PC2 (9.61% variance explained) underscored stark differences in total transcription signatures between brain tissue and the other tissues. When examining the differential gene expression between the salt-polluted and the reference stations, significant numbers of genes were identified for each tissue. In gills, we found 286 DEGs (117 down- and 170 up-regulated) in the salt-polluted compared to the reference station **(Figure 2B).** For brain tissue, 1350 DEGs (743 down- and 605 up-regulated) were observed in the salt-polluted compared to the reference **(Figure 2C).** In the liver, we found 673 DEGs (426 down- and 248 up-regulated) in the salt-polluted compared to the reference **(Figure 2D).** Notably, across all three tissue DEGs, the pattern of expression was consistent among the three sample replicates per station.

**Figure 2.**
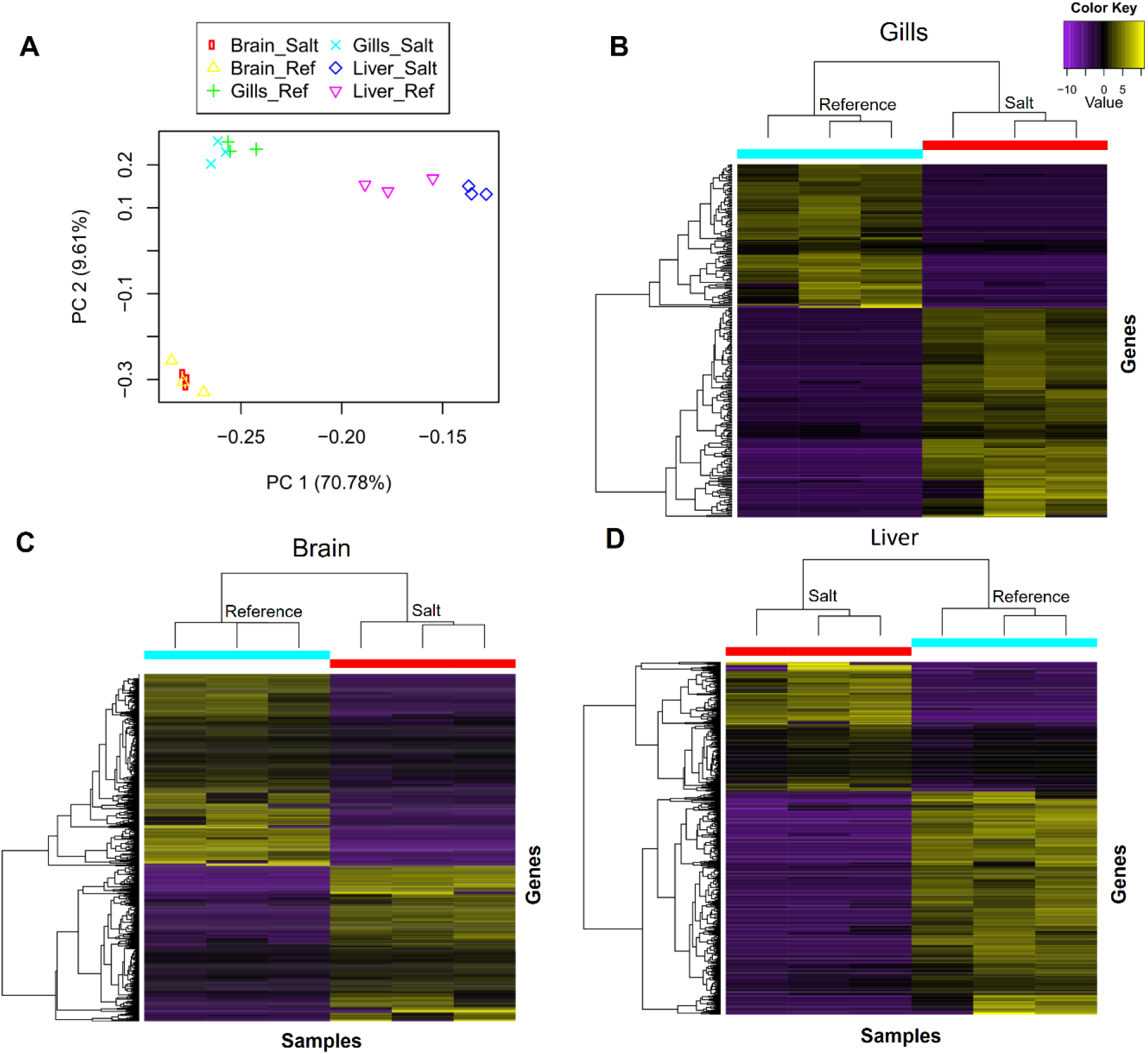
**A.** Principal component analysis (PCA) of gills, brain, and liver transcriptomes of minnow (*Phoxinus septimaniae* x *P. dragarum*) sampled at a salt-polluted (Salt) and a reference station (Reference).**B.** Heatmap displaying 286 differentially expressed genes (DEGs) in gills between the salt and Reference stations. **C.** Heatmap displaying 1350 differentially expressed genes (DEGs) in the brain between the salt and Reference stations. **D.** Heatmap displaying 673 differentially expressed genes (DEGs) in the brain between the salt and Reference stations. For all heatmaps, the X-axis represents sample replicates per station, and the Y-axis represents individual gene expression. Upregulated genes are shown in yellow, with brighter colors indicating higher expression values. In contrast, purple shades indicate downregulated genes, with the brightest shade indicating the strongest downregulation.

The significant DEGs for all of three tissues were assigned to three major Gene Ontology (GO) categories: biological process, cellular component, and molecular function. The DEGs were classified in a total of 147, 197 and 182 higher-level GO terms relative to the reference station for the gills, brain and liver, respectively (**Table S1**). Interestingly, when the higher-level gene ontology terms were analyzed with a Venn diagram for all three tissues we found a significant number of terms shared with all tissues **(Figure 3).** In the biological process category, 68 terms were shared among all three tissues (**Figure 3**). In the cellular component category, 24 terms were common, and in the molecular function category, a total of 26 terms were shared across all tissues.

**Figure 3.**
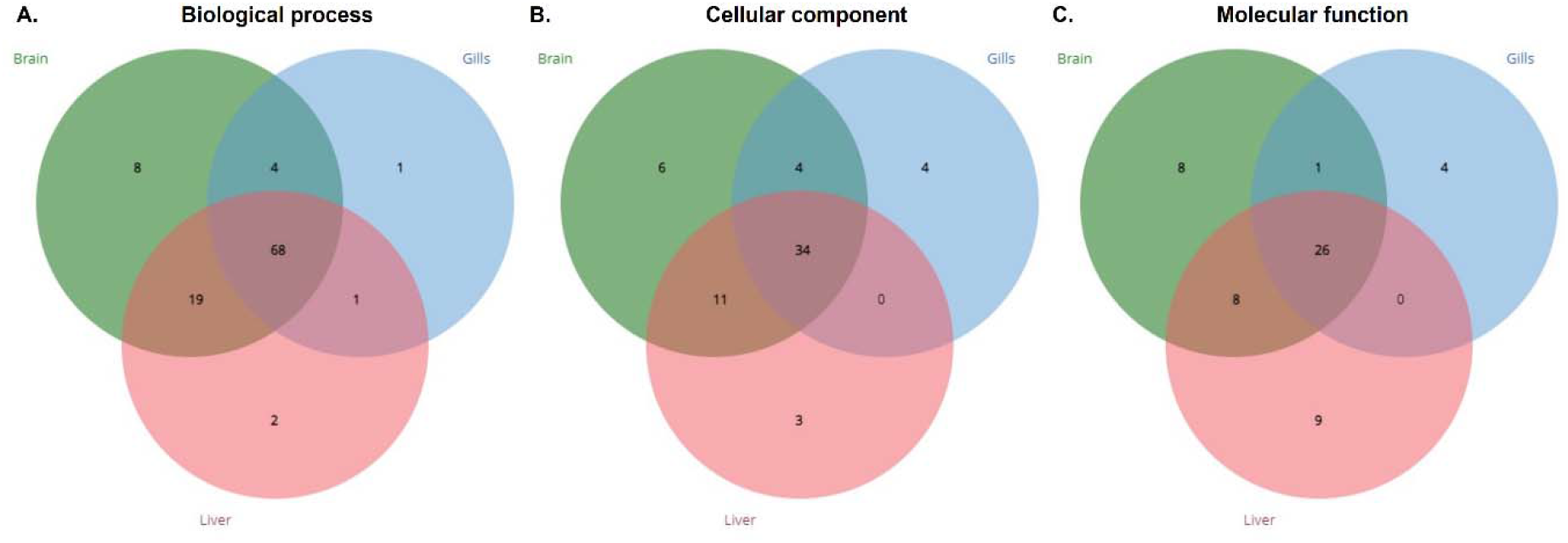
Venn diagrams illustrating shared go higher-level terms across tissues. A. 68 terms shared in the Biological Process category. B. 34 terms shared in the Cellular Component category. C. 26 terms shared in the Molecular Function category. Overlap of gene ontology terms is represented by the intersection of circles, each corresponding to brain (green), liver (red), and gills (blue).

Key terms related to osmotic and chemical stress were graphically presented, showing the shared and distinct terms among the three tissues (**Figure 4**). Noteworthy terms included hydrolase activity, small molecule binding, carbohydrate derivative binding, transporter activity, transmembrane transporter activity, and channel regulator activity in the molecular function category. In the cellular component category, terms such as organelle membrane, cell junction, extracellular region, and synapse were common across all tissues. In the biological process category, significant terms included responses to chemical stress, catabolic processes, developmental process regulation, signaling, immune system processes, growth, reproduction, and methylation, all shared across the three tissues.

**Figure 4.**
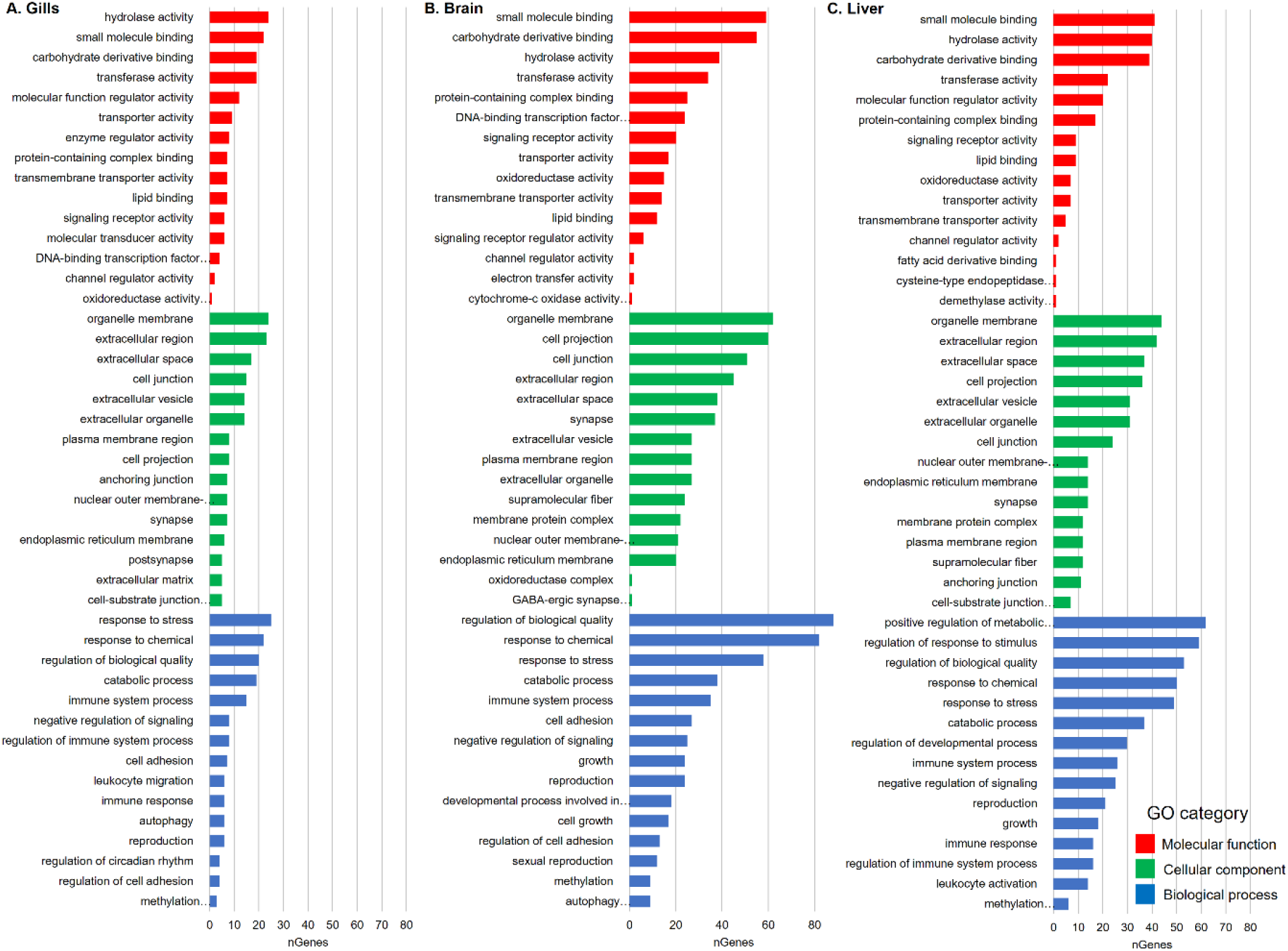
Functional classification of differentially expressed genes in the high-level GO pathway categories across brain, liver, and gill tissues. The Y-axis represents the different GO general categories, with different colors for each one. And the X-axis represents the number of genes (nGenes) that were annotated to each pathway.

The complete set of genes that were found to be differentially regulated for all tissues in the salt station compared to the reference were included in the enrichment analysis **(Table S2).** The top 10 significantly enriched pathways and those found to be related to salinity and overall chemical stress for the different tissues are presented graphically in **Figure 5**. In gills for the molecular function category, the most significant pathway was the “chloride channel inhibitor activity”, constituted by two genes *WNK2* and *CFTR* **(Figure 5A).** These two genes appeared as constituents of several other ion transport and channel regulation-related pathways annotated in the enrichment analysis together with genes like *KCNC, ANXA2, ANO10, and CUL5* **(Table S2).** In the cellular component category, the “endomembrane system” pathway had the highest number of genes, followed by “chromosome, telomeric region”, and pathways related to the vacuole and lysosome. In the biological process the pathways with the highest number of genes and higher significance were the “cellular response to stimulus”, and “cellular response to stress”, followed by pathways related to the transport of lipids.

**Figure 5.**
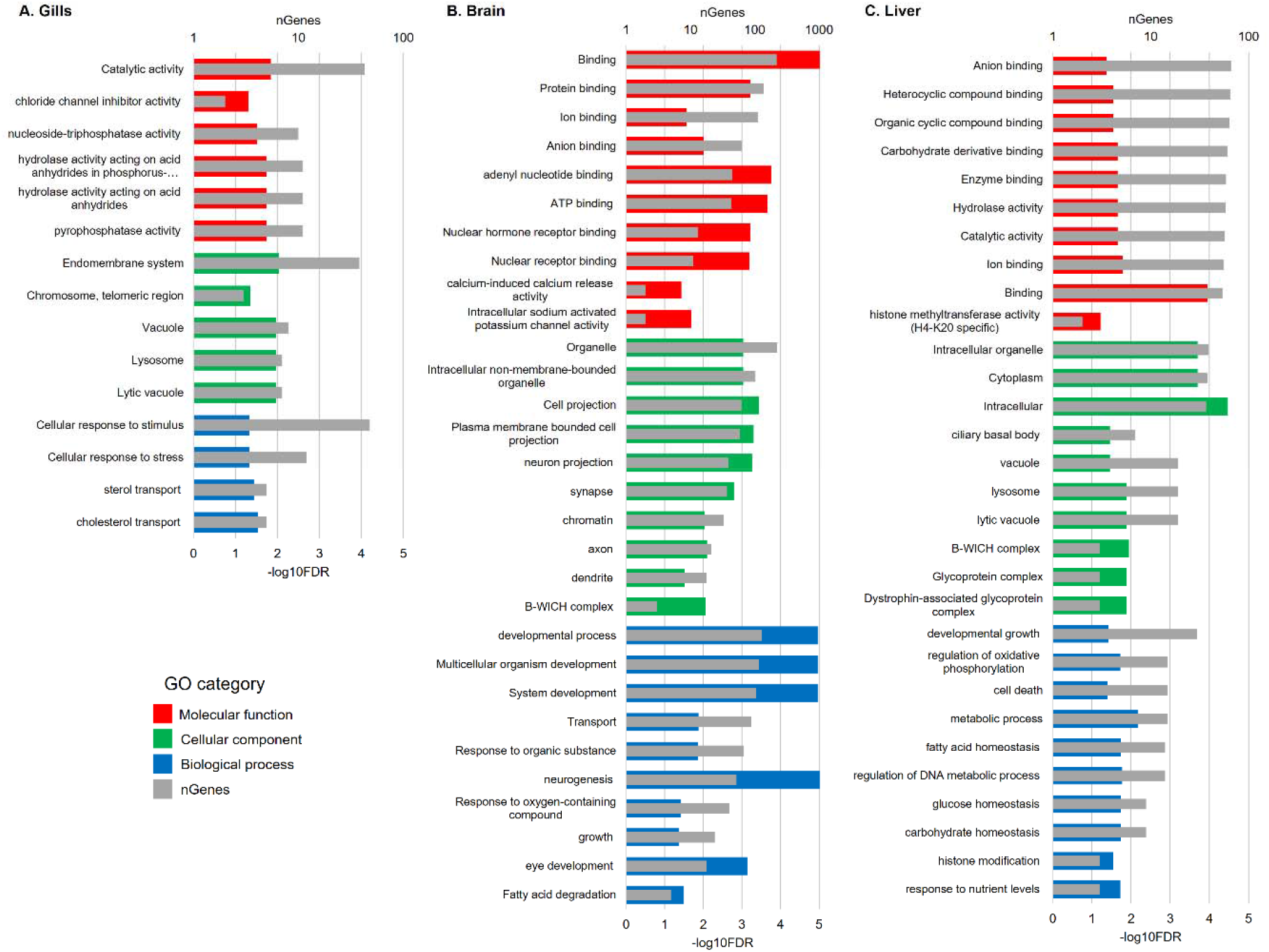
Pathway enrichment analysis showing the top 10 enriched pathways related to salt or chemical stress across gills, brain, and liver tissues. The superior X-axis represents the number of genes annotated in the pathway (nGenes) and is expressed in log10 for the sake of the comparison between tissues. The inferior X axis is the enrichment FDR and is represented in – log10. The Y-axis is the top 10 enriched pathways related to salt or chemical stress. A. represents the gill-enriched pathways. B. the brain pathways and C. the liver.

In the brain, for the molecular function category, the pathway with the highest number of genes and more enrichment significance was “Binding” followed by other pathways related to Protein, Ion, Anion, and ATP binding **(Figure 5B)**. It is worth noting that several other pathways with less significance that were related to channel activity and overall transport were recurrent in the enrichment analysis with representative genes as *CLCN2, ANO1, KCNN3, KCNT1, KCNT2, RYR2, RYR3* **(Table S2).** For the cellular component, the “organelle” pathway presented the most genes, followed by “cell projection”, other pathways related to neuronal activity like “neuron projection”, “synapse and “axon” were significantly enriched. For the biological process the most significant enriched pathways were those related to development, followed by the “transport” pathway, and the response to organic substances and oxygen-containing compound. Interestingly the “growth”, “neurogenesis” and “eye development” pathways were also enriched in the brain. A detailed examination of the differentially expressed genes in the brain revealed immune system-related genes such as *RPTOR* and *MKNK2*. It is worth noting that the “Fatty acid degradation” pathway was the only KEGG pathway that was found to be enriched in the analysis and it is represented in **Figure 6**. In the figure, the red highlighted boxes represent the genes found in this study: *CPT1A*, *ACSL1* (6.1.2.3), *ACSL4* (6.1.2.3), *ACADM* (1.3.8.7) *ACADS* (1.3.8.1).

**Figure 6.**
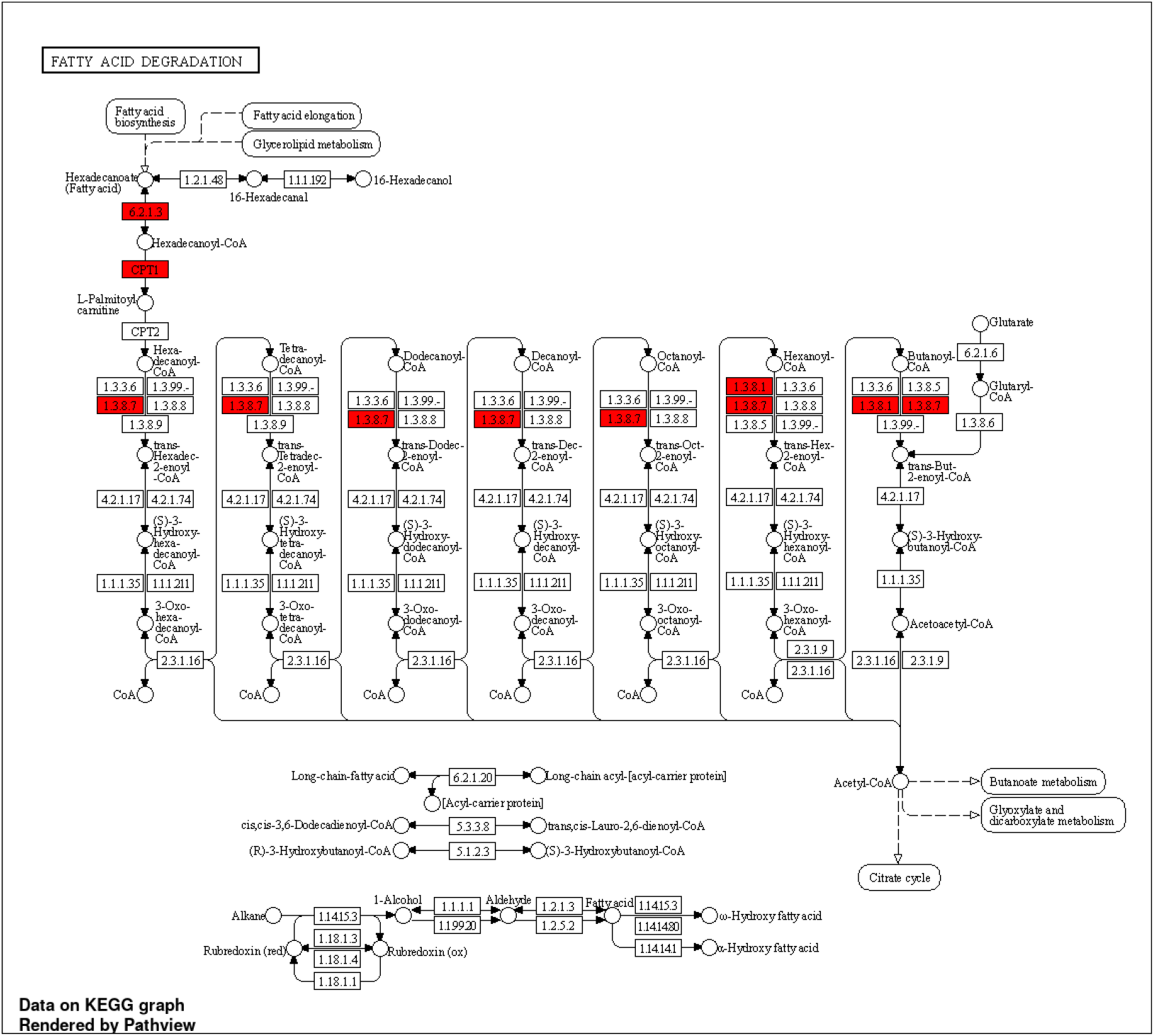
KEGG enriched pathway "Fatty acid degradation" for brain tissue. Red highlighted boxes represent differentially expressed genes when comparing the salt and reference stations.

In the liver similar to our findings in the brain, for the molecular function category, the most important enriched pathways were related to binding, “anion binding” the one with more genes **(Figure 5C)**. In this category are also noteworthy the enrichment of the hydrolase and catalytic activity pathways. In the cellular component category, the more enriched pathways are the intracellular organelle and the cytoplasm. The enriched pathways “vacuole”, “lysosome” and “lytic vacuole” are recurrent in the liver as it was in the gills. And the “B-WICH” complex enrichment is shared with the brain. For the biological process category, the most enriched pathway was the “developmental growth”, followed by the “regulation of oxidative phosphorylation” and “cell death”. An enrichment of metabolic-related pathways was also evident with pathways such as “metabolic process”, fatty acid, glucose, and carbohydrate homeostasis. Also, it is remarkable the enrichment of the “histone modification” and “response to nutrient levels”. Additionally, the liver exhibited significant differential expression integral to immune-related pathways, notably including heat shock proteins (HSPs) such as HSP70, HSP7C, HSP7E, and the heat shock transcription factor 2 (HSF2).

### 2.3. RNA-seq data validation by qPCR

Validation of the transcriptomic data for the 24 chosen genes using qPCR was conducted using a Spearman correlation analysis of the log2 fold changes for the 18 samples analyzed on both platforms. Overall, there was a significant correlation between the log2 fold changes between platforms (Spearman, R = 0.71, p < 0.001; **Figure 7**)

**Figure 7.**
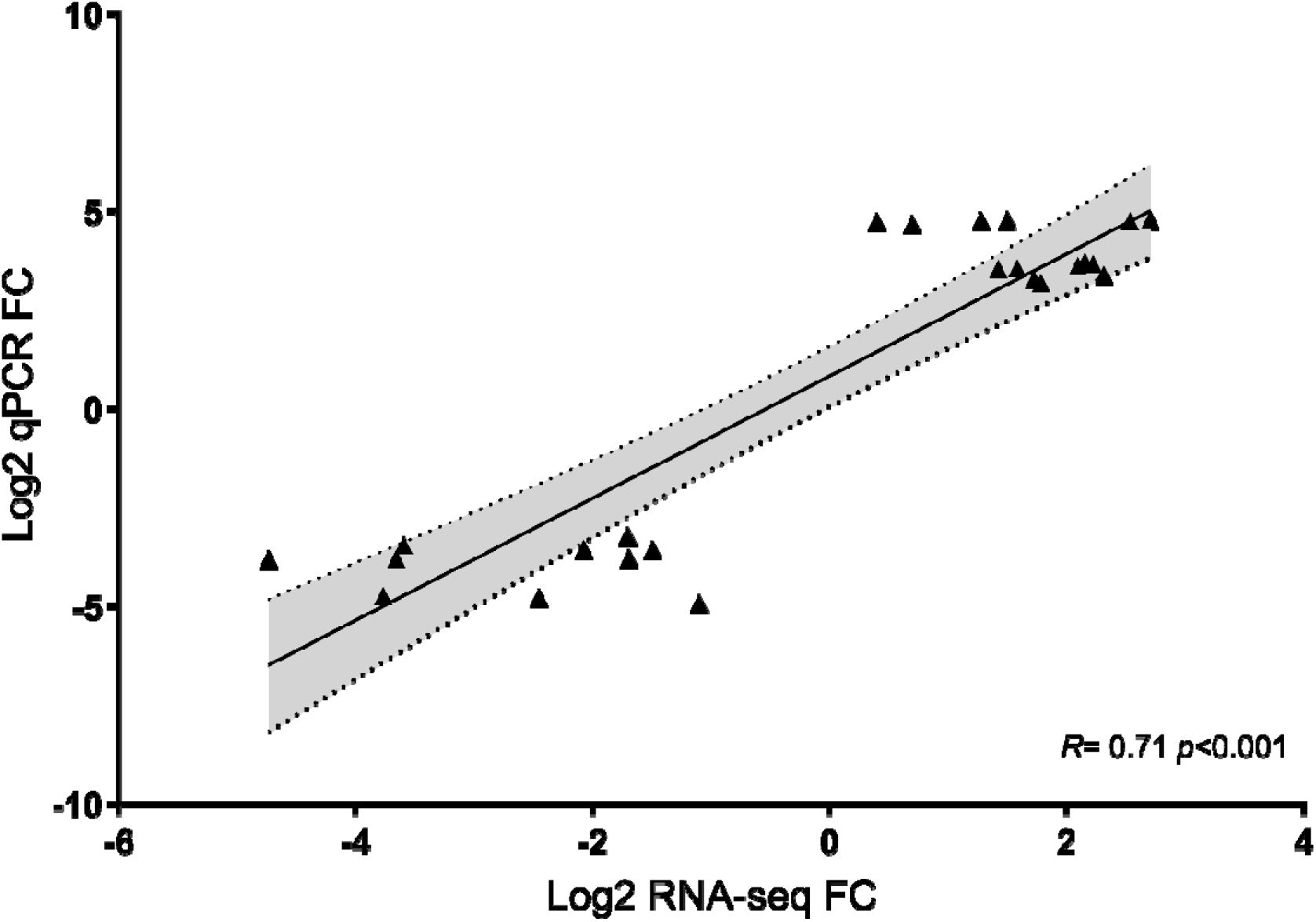
Validation of RNA-seq data with quantitative PCR (qPCR) for the 24 genes analyzed on both platforms. Log2 fold changes for each gene are plotted, and the relationship was analyzed using a Spearman correlation. The reference line indicates a linear relationship between the RNA-seq and qPCR results. The shaded area represents a 95% confidence interval.

## 3. Discussion

In summary, the study conducted extensive RNA-seq analysis on the brain, gill, and liver tissues of the minnow across two distinct salinity stations, revealing profound alterations in gene expression profiles induced by severe salinity stress. Notably, the differential gene expression patterns between the *s*alt-polluted and reference stations demonstrated substantial differences across tissues, with the brain showing the highest number of differentially expressed genes. The GO and pathway enrichment analyses unveiled shared biological processes, molecular functions, and cellular components, emphasizing the impact of osmotic and chemical stress in all tissues. Specifically, the brain, gills, and liver exhibited tissue-specific responses to salinity changes, illuminating the intricate molecular adaptations essential for osmoregulation and stress responses. The validation of RNA-seq data through qPCR reinforced the robustness of the findings. Furthermore, The high quality of raw data and quality control results of our transcriptome, together with a mapping rate for all tissues on par with other studies (Guo et al., 2018b; Sun et al., 2020), demonstrate the effectiveness and reliability of the high-throughput sequencing transcriptomic analysis.

The overall differential expression of pathways, including responses to chemical stress, catabolic processes, developmental process regulation, signaling, immune system processes, growth, reproduction, and methylation, shared between tissues serves as an indicator of stress originating from salinity and the multiple stressors found in the Llobregat River. From the various enriched pathways previously cited we highlight the ones related to differential expression of growth and reproduction which previous transcriptomic studies have linked to long-term salinity stress (K. Zhou et al., 2020). Another set of relevant pathways are those related to neurogenesis and eye development have been related to the effect multiple chemical stressors in previous transcriptomic studies (Meade et al., 2023), and in our case can be related to the multiple stressors that act together with salt pollution in the Llobregat River. Interestingly, the "methylation" pathway was enriched in all tissues, and in the liver, "histone modification," as suggested by a recent study coupling methylation and transcriptomics in a euryhaline fish, indicates methylation changes and gene expression patterns under salinity stress (Blondeau[Bidet et al., 2023). As we are investigating the transcriptomic response of three tissues in a field setting where multiple stressors act simultaneously, and the complexity of the enriched metabolic pathways and their interplay increases, a thorough discussion of all enriched pathways would be beyond the scope of this study. We therefore focused the discussion to osmoregulation, metabolic pathways, and immune responses, recognizing them as the primary pathways affected by our dominant salt pollution stressor.

### 3.1. Osmoregulation

We identified differentially expressed pathways related to osmotic stress and general chemical stress in all three gene ontology categories shared between all tissues under the salt-polluted station compared to the reference. We focused our discussion on terms related to salinity and chemical stress. Notably, in the molecular function category, pathways such as transporter activity, transmembrane transporter activity, and channel regulator activity were of particular importance. These terms comprise proteins responsible for moving ions across membranes, ion channels, ion pumps, and ion transporters essential for maintaining osmotic pressure in exchange for considerable energy (Huang et al., 2004). Studies on the gills of various fish species in varying saline habitats have shown a significant number of differentially expressed genes (DEGs) involved in ion transport and transmembrane transport (X. Chen et al., 2021; Escobar-Sierra & Lampert, 2023; Gibbons et al., 2017; Guo et al., 2018b; Su et al., 2020a). Specifically, for gill tissue, we found enrichment of osmoregulation-related pathways like “chloride channel inhibitor activity”. Genes constituting the main channel regulation-related pathways for the gills included WNK2, CFTR, KCNC, ANXA2, ANO10, and CUL5. Anoctamins, a family of Ca^2+^-activated Cl^−^ channels and phospholipid scramblases, have been shown to support cell volume regulation (Hammer et al., 2015). WNK2 regulates sodium-coupled chloride cotransporters and is part of an essential pathway for regulating cell volume in response to osmotic stress (Kahle et al., 2010). ANXA2 is involved in cellular differentiation and ion channel conductance, with its differential expression recorded in euryhaline fish (Boulet et al., 2012). KCNC, like other voltage-gated Ca2+ channels, has been demonstrated to be involved in processes pertinent to the regulation of ion and water transport in animal epithelia— Ca^2+^ signaling and entry regulation (Zheng & Trudeau, 2015). The cystic fibrosis transmembrane conductance regulator (CFTR) is expressed on the apical surface of branchial ionocytes, serving as the conduit for active Cl^−^ secretion and an ion channel regulator in euryhaline fish (Shaughnessy & Breves, 2021). *CUL5* is a protein that mainly controls osmoregulation and regulates blood pressure within the body (Xu et al., 2021).

While the brain has not been traditionally studied in relation to osmoregulatory stress, its high importance in osmotic homeostasis has been reported, with the enrichment of osmoregulatory-related pathways reported under salinity stress in fish (Liu et al., 2018). In our study, the brain exhibited transport enrichment and overall osmoregulation, evident through the differential expression of genes like *CLCN2, ANO1, KCNN3, KCNT1, KCNT2, RYR2*, and *RYR3*. Chloride channels play a key role in the osmoregulatory physiology of all animals, with CLCN2 differentially expressed in euryhaline fish exposed to osmotic stress (Bonzi et al., 2021; Su et al., 2020b). Another member of the anoctamin family, *ANO1*, related to osmoregulation, was differentially expressed in fish under osmotic stress (Taugbøl et al., 2022). *KCNN3* is a potassium-activated channel that has been proven to be differentially expressed in euryhaline fish exposed to osmotic stress (Bonzi et al., 2021). The potassium channels *KCNT1* and *KCNT2* are activated under high concentrations of chloride and be differentially expressed in euryhaline fish exposed to osmotic stress (Blondeau[Bidet et al., 2023; Lüscher et al., 2020). *RYR2* and *RYR3* are known interaction partners in the formation of chloride channels that are key in several physiological processes, including osmoregulation (Zeng et al., 2018). In summary, our results suggest that salinity from potash mining in the Llobregat is influencing the activation of pathways related to osmoregulatory stress and homeostasis in both gills and brain tissue. Furthermore, we were able to pinpoint several candidate genes related to these pathways that could serve as biomarkers of this type of stress for further studies.

### 3.2. Metabolism

In all three tissues, we observed differential gene expression related to metabolism, particularly notable in the liver, where various metabolic pathways were activated, and in the brain, with enrichment in the KEGG pathway ’Fatty acid degradation.’ Osmoregulatory processes inherently demand energy, and the roles of lipid, glucose, and carbohydrate metabolism in fish osmoregulation have been extensively reviewed (Tseng & Hwang, 2008). Fish increase lipid metabolism primarily in the liver and brain during salinity stress, as reported in other transcriptomic studies of species facing hypersaline conditions (Gibbons et al., 2017; Hu et al., 2015). Specifically in the liver, enrichment of lipid metabolism under hypersaline stress for a euryhaline fish has included pathways such as fatty acid elongation, fatty acid metabolism, and fatty acid biosynthesis (K. Zhou et al., 2020). Both fish brain and liver tissues have shown an enrichment of pathways related to glucose and lipid metabolism when challenged with a strong salinity gradient (Hu et al., 2015). Furthermore, some genes differentially expressed under osmotic stress in the brain, involved in the fatty acid degradation pathway, have been previously reported in studies of species undergoing osmoregulation stress. For instance, carnitine palmitoyltransferase 1 (*CPT1*) and overall lipid catabolism were found to be differentially expressed under osmoregulatory stress in the marine euryhaline crab *Scylla paramamosain* (Luo et al., 2023). *ACAD* genes, encoding acyl-CoA dehydrogenases, were differentially expressed under osmoregulatory stress in *Eriocheir sinensis*, an extremely invasive alien crab species (Hui et al., 2014). The enrichment of the fatty acid degradation pathway in the brain and other metabolic-related pathways in the liver in our study indicates that the minnow activates metabolic pathways to meet the energy demands associated with physiological adaptations required to survive hypersaline stress.

### 3.3. Immune and stress response

We observed an overall differential expression of immune system processes and chemical stress across all tissues in the salt-polluted station. The relationship between immunity and stress has been extensively reviewed, suggesting that long-term exposure to salinity and other chemical stressors significantly influences fish immune responses (Tort, 2011; Z. Zhou et al., 2021). In line with this, we identified pathway enrichment and differentially expressed genes associated with stress and immune system regulation. A notable finding was the significant differential regulation of a set of Heat shock proteins (HSPs) in the liver tissue, including HSP70, HSP7C, HSP7E, and the heat shock transcription factor 2 (HSF2). These HSPs play a crucial role in translocation and protein folding, commonly serving as indicators and biomarkers of abiotic stress, expressed under salinity stress in several fish species (Jeffrey et al., 2023; Li et al., 2020; Lin et al., 2020; Puntila-Dodd et al., 2021). Another indicator of immune system alteration was the differential expression of the genes RPTOR and MKNK2 in the brain tissue under salinity stress. The MAPK interacting serine/threonine kinase 2 (MKNK2) is a substrate of the MAPK pathway, crucial in its regulation (Maimon et al., 2014). The Mitogen-activated protein kinase (MAPK) signaling is known to be involved in various cell processes, including innate immunity (Roux & Blenis, 2004). Furthermore, its differential expression has been reported and linked to fish immune response when challenged with chemical stressors, including salinity (Duan et al., 2022; Tian et al., 2019). The regulatory-associated protein of MTOR complex 1 (*RPTOR*) is associated with the mammalian target of rapamycin (mTOR), which plays a key role in controlling and shaping the effector responses of innate immune cells (Weichhart et al., 2015). The activation of this pathway has been identified using transcriptomics in the euryhaline fish *Oreochromis mossambicus* following osmotic stress (Su et al., 2023). In conclusion, our findings highlight a comprehensive differential expression of immune system processes and stress-related pathways, emphasizing the intricate molecular responses of minnow to salinity stress in a polluted environment.

### 3.4. Ecological implications

Although our study design does not directly measure other stressors known to affect the Llobregat River, such as sewage discharges and emergent pollutants altering fish fauna (Munné et al., 2012), our transcriptomic results enable us to infer physiological responses to the known severe changes in salinity and unknown multiple stressors in this highly polluted urban river. At our study stations, salinity pollution from potash mining emerged as a dominant stressor, potentially creating a bottleneck for sensitive, freshwater-adapted native fishes.

Minnows in the highly polluted Llobregat exhibit physiological adaptations to significant changes in salinity and various anthropogenic stressors. Our study unveiled the activation of multiple physiological pathways, evidence for the species’ ability to survive despite potentially high physiological costs. This adaptability aligns with traits associated with invasiveness and the high adaptive potential of other invasive species in the face of new stressors (Davidson et al., 2011). In fact, the minnow’s invasion success in the highly polluted Llobregat is documented by their three-fold increase in occurrence in the catchment between the 1990s and 2003, while the individual number and species diversity of native fish declined sharply at the same time (Maceda[Veiga et al., 2010).

Our transcriptomic approach provides insights into the molecular pathways activated by fish in response to the complex challenges posed by a heavily polluted river affected by multiple anthropogenic stressors. These pressures, resulting from human interventions, act as adaptive forces by creating distinct environmental conditions and pressures that select invasive species and specific traits in those species (Cadotte et al., 2017). For instance, invasive fish species with increased thermal tolerance, salinity tolerance, and physiological adaptability—traits that enhance tolerance to urban environments—are prevalent in urban areas (Gomes-Silva et al., 2020; Green et al., 2023). This supports the proposition of Borden & Flory, (2021), which suggests that urban areas create evolutionary pressures altering invasive species adaptations, increasing their potential to rapidly spread and succeed over native species. Our results highlight the detrimental impacts of current practices in potash mining, urban expansion, and overall anthropogenic interventions on European freshwater fauna. In light of our research, we propose that management strategies for controlling the invasion of *Phoxinus septimaniae* x *P. dragarum* in the Llobregat catchment should prioritize efforts to mitigate pollution originating from potash mining. Addressing this specific source of stress is crucial to curbing the physiological adaptive advantage conferred to invasive species over their native counterparts.

## 4. Conclusion

In summary, our investigation delved into the intricate molecular mechanisms governing freshwater salinization adaptation in invasive minnow populations within the Llobregat River, Barcelona, Spain. This study shed light on the comprehensive alterations in global gene expression profiles across the brain, gills, and liver tissues in response to contrasting salinity levels observed between the salt-polluted station and reference stations. Notably, the brain displayed the most distinct sensitivity to salinity, with the highest number of differentially expressed genes, emphasizing its vital role in salinity adaptation. Tissue-specific pathways related to stress, reproduction, growth, immune system responses, methylation, and neurological development were identified, marking their relevance in the context of heightened salinity stress. Through the rigorous validation of our RNA-seq data using quantitative PCR, we reinforced the reliability and consistency of our findings across platforms.

Our exploration, unveiling shared and distinct molecular responses, contributes significantly to our understanding of the genetic and physiological mechanisms pivotal for fish adaptation in salinity-stressed environments. These findings align with and further clarify previous studies on fish transcriptomics, supporting and extending existing knowledge in this field. Additionally, we were able to measure a physiological response to a stressor in wild non-model organisms within field conditions, where multiple stressors may interact. This aspect emphasizes the importance of our study. In line with our prior research, we highlight the potential of transcriptomic approaches in evaluating the physiological responses of wild organisms facing known or unknown stressors. This capability bolsters our assertion that transcriptomics is essential for guiding management and policy decisions, providing valuable insights into the complex interactions of diverse stressors in natural settings and enabling more informed conservation strategies.

The analysis of tissue-specific pathways and shared biological processes elucidates the complexity of osmoregulation and stress responses in aquatic organisms facing freshwater salinization. The results, presenting a detailed transcriptomic profile of adaptation, provide crucial insights for the conservation and management of freshwater ecosystems challenged by increasing salinity pressure. This study opens new avenues for further research and forms a foundational basis for addressing the multifaceted impacts of salinity stress.

## 5. Acknowledgements

This paper benefited from the multiple discussions within the Collaborative Research Centre 1439 RESIST (Multilevel Response to Stressor Increase and Decrease in Stream Ecosystems; www.sfb-resist.de). Funding was provided by the Deutsche Forschungsgemeinschaft (DFG, German Research Foundation; CRC 1439/1, project number: 426547801).We thank the Regional Computing Center of the University of Cologne (RRZK) for providing support and computing time on the High high-performance computing (HPC) system CHEOPS. And finally, we thank the Cologne Center for Genomics (CCG), for their technical support on sample preparation and sequencing.

## Notes

### Competing Interest Statement

The authors have declared no competing interest.

